# Shared representations of human actions across vision and language

**DOI:** 10.1101/2023.11.06.565690

**Authors:** Diana C. Dima, Sugitha Janarthanan, Jody C. Culham, Yalda Mohsenzadeh

## Abstract

Humans can recognize and communicate about many actions performed by others. How are actions organized in the mind, and is this organization shared across vision and language? We collected similarity judgments of human actions depicted through naturalistic videos and sentences, and tested four models of action categorization, defining actions at different levels of abstraction ranging from specific (action verb) to broad (action target: whether an action is directed towards an object, another person, or the self). The similarity judgments reflected a shared organization of action representations across videos and sentences, determined mainly by the target of actions, even after accounting for other semantic features. Language model embeddings predicted the behavioral similarity of action videos and sentences, and captured information about the target of actions alongside unique semantic information. Together, our results show how action concepts are organized in the human mind and in large language model representations.

## Introduction

Daily life requires us to understand a wide range of actions performed by others, as well as communicate about them. Yet actions are an incredibly heterogeneous category, linking multiple visual, semantic and social domains^1–4^ and varying along multiple axes. How does the mind organize this rich semantic space, and does this organization differ when actions are communicated visually or verbally?

Actions can be defined at multiple levels of abstraction and can only be understood in relation to agents, objects, and contexts, a complexity that still challenges machine learning algorithms for action recognition. Different brain areas have been shown to represent action categories with different degrees of specificity^5–7^. Despite this, previous work on semantic action representations has tended to either define action categories *a priori* (e.g. using action verbs or broad activity types)^8–10^, or to employ a behaviorally estimated semantic space^11,12^. Recent work has investigated the taxonomy of actions using text stimuli of varying levels of abstraction^13^; but it is unclear whether such a taxonomy would generalize to visual actions and different categorization tasks.

Behavioral studies using visual actions and data-driven analyses have mapped representational spaces along broad axes such as *communication*, *locomotion*, *food* etc.^11,14,15^ or axes related to facial traits, emotions, and action planning^16^. In a large-scale text analysis, six dimensions organized action representations^17^, some of them notably more abstract than their visual counterparts (e.g. *abstraction*, *tradition*, *spirituality*). In light of such results, it has been speculated that the dimensions organizing actions in the mind may be modality-dependent^14^; but to our knowledge, no behavioral study to date has investigated semantic action representations using both visual and text stimuli.

However, a growing body of neuroimaging research suggests that action representations may share a neural substrate across vision and language. Adding to long-standing evidence of overlap in neural activations elicited by action images and action verbs^10,18–24^, recent investigations have more directly revealed modality-invariant neural representations in human occipito-temporal and parietal cortices^25,26^. These areas have also been shown to encode invariant action category^1,11,27–33^, hinting at a putative cross-modal semantic architecture.

Here, we employed semantic similarity to capture how action categories are organized in the mind^34–36^. To capture a naturalistic action space^37–39^, we curated a set of videos and sentences depicting everyday actions defined at four levels of abstraction based on prior research: specific action verbs^10,26^ (e.g. *swimming*), everyday activities^8,9,40^ (e.g. *sports*), broad action classes inspired by primate research^41,42^ (e.g. *locomotion*), and action target types, reflecting whether an action is directed towards an object, another person, or the self^12,43^.

To understand how actions are organized in the mind, we used these semantic features, together with a rich set of contextual, perceptual, and computational features, to predict behavioral similarity judgments of the videos and sentences within and across modalities. We found a semantic organization shared across vision and language, reflecting the target of actions and, to a lesser extent, their class. Pre-trained text embeddings based on OpenAI’s GPT large language models (LLM) predicted this shared semantic structure better than, and independently of, other semantic features, while also containing information about the target of actions. Together, these results reveal the modality-invariant organization of action concepts in minds and machines.

## Results

### Features in naturalistic actions: from semantic to perceptual

We curated a set of 95 videos of natural actions from the Moments in Time dataset^44^. Videos were carefully selected to ensure that semantic, contextual, and perceptual features were minimally correlated across the dataset (**Figure 1b**). For each video, a corresponding naturalistic sentence was constructed following the structure *agent* + *action* (+ *object* if present) + *context*, e.g. *Women are swimming in a pool*. The sentences were displayed as images matched in size to the videos (see Methods: Stimuli).

**Figure 1.**
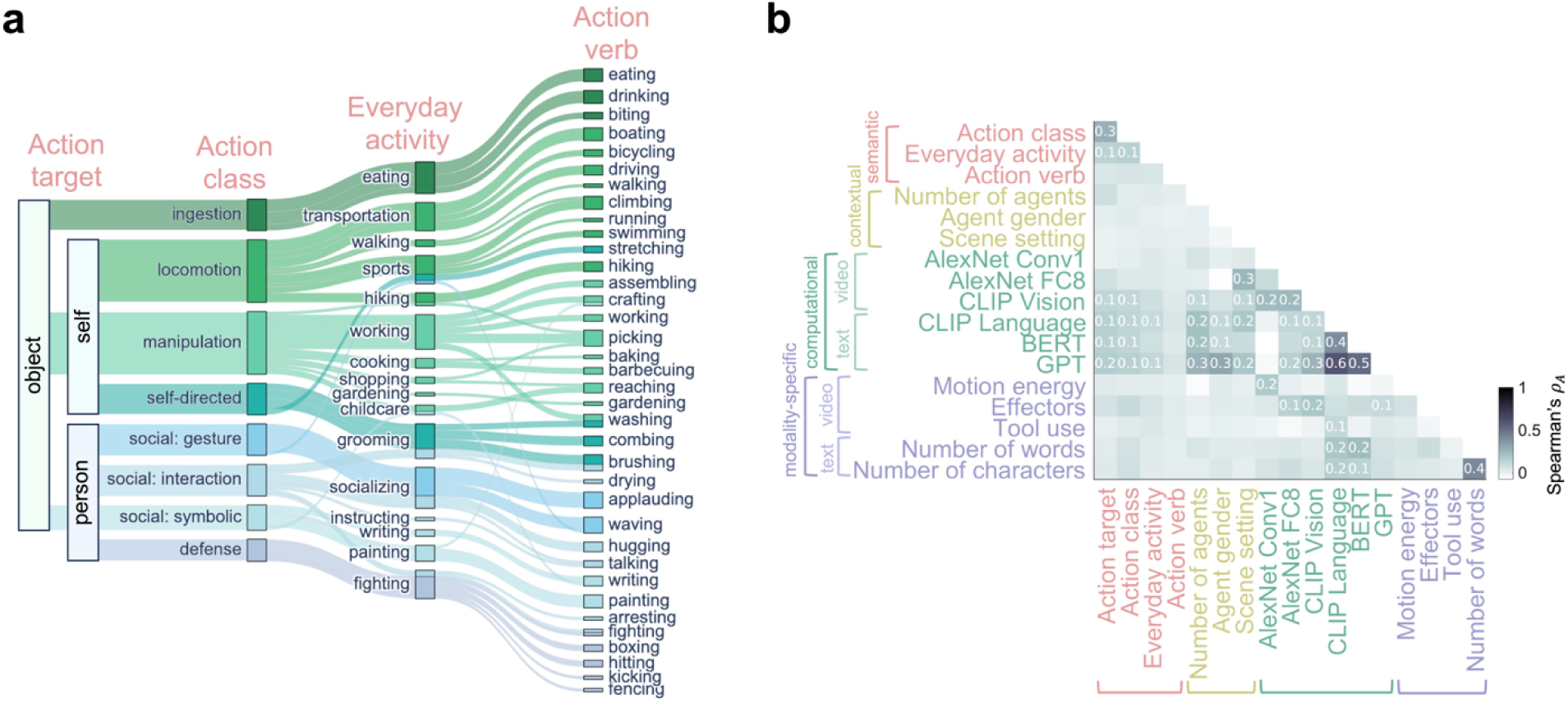
Action features across the stimulus set. **a,** Actions were categorized at different levels of abstraction, from specific to broad, according to their action verb, everyday activity category, action class, an action target. **b,** Correlations across features of interest were low, with the notable exception of more comple computational models (CLIP, BERT, GPT embeddings) that moderately correlated with most other features. Both video- and text-based feature representational dissimilarity matrices (RDMs) are included in the correlation matrix, and only correlations over 0.1 are displayed numerically.

We labeled the stimuli with a set of 18 semantic, contextual, computational, and modality-specific features. First, actions were labeled with four semantic features at different levels of abstraction based on previous research (**Figure 1a**): by their action verb^10,26,45^ (e.g. *swimming*), everyday activity^15,46^ (e.g. *sports*), action class^41^ (e.g. *locomotion*), and action target^12,43^ (e.g. *the self*). The first three roughly correspond to the subordinate, basic and superordinate levels of categorization identified in object research^47–49^, while at the same time testing the main action definition criteria prevalent in the literature. The fourth feature (action target) includes information about the actions’ goals and their degree of sociality (person-directedenss) and transitivity (object-directedness), features previously found to be essential in action processing^9,12,43^.

Apart from their semantic content, naturalistic actions vary along numerous axes, including those related to the agents performing the action and the context in which the action takes place. To capture this information, we labeled each action with contextual features: the number and perceived gender of agents, as well as the scene setting (indoors or outdoors).

Next, to better characterize stimulus features and evaluate how well computational models can capture human action representations, we extracted image features from different vision-trained neural networks (the first convolutional and last fully connected layers of AlexNet^50^, the vision module of CLIP^51^) and sentence features based on large language models (LLM: CLIP’s language module, BERT^52^, OpenAI GPT text embeddings^53^).

Finally, we accounted for additional sources of variance by extracting modality-specific features encoding perceptual and action information: motion energy, effectors (body parts involved in each action), and tool use (for videos) and the number of words and characters (for sentences).

We generated representational dissimilarity matrices (RDMs) quantifying the pairwise distances between all videos or sentences along each of the 18 semantic, contextual, computational, and modality-specific features (see Methods: Feature definitions).

Correlations among different groups of features were low across both videos and sentences (**Figure 1b**). There were moderate correlations among semantic features, particularly between action target and class (p_A_=0.27). Features extracted from computational models (particularly LLMs) moderately correlated with semantic, contextual, and perceptual features, pointing to the rich, diverse information captured by these models.

### A shared organization across vision and language

To investigate how humans represent the meaning of actions, we conducted two multiple arrangement experiments with either the video or sentence stimuli. Participants arranged the videos or sentences inside a circular arena (**Figure 2a**) and were instructed to place similar actions closer together and different actions further apart, focusing on the meaning of the actions. Different sets of 3-12 videos were displayed during the task, until all pairwise distances were reliably estimated. This method efficiently captures multi-dimensional representations by presenting the stimuli multiple times in different contexts. On-screen distances were used to generate behavioral RDMs for each participant using inverse MDS^54^.

**Figure 2.**
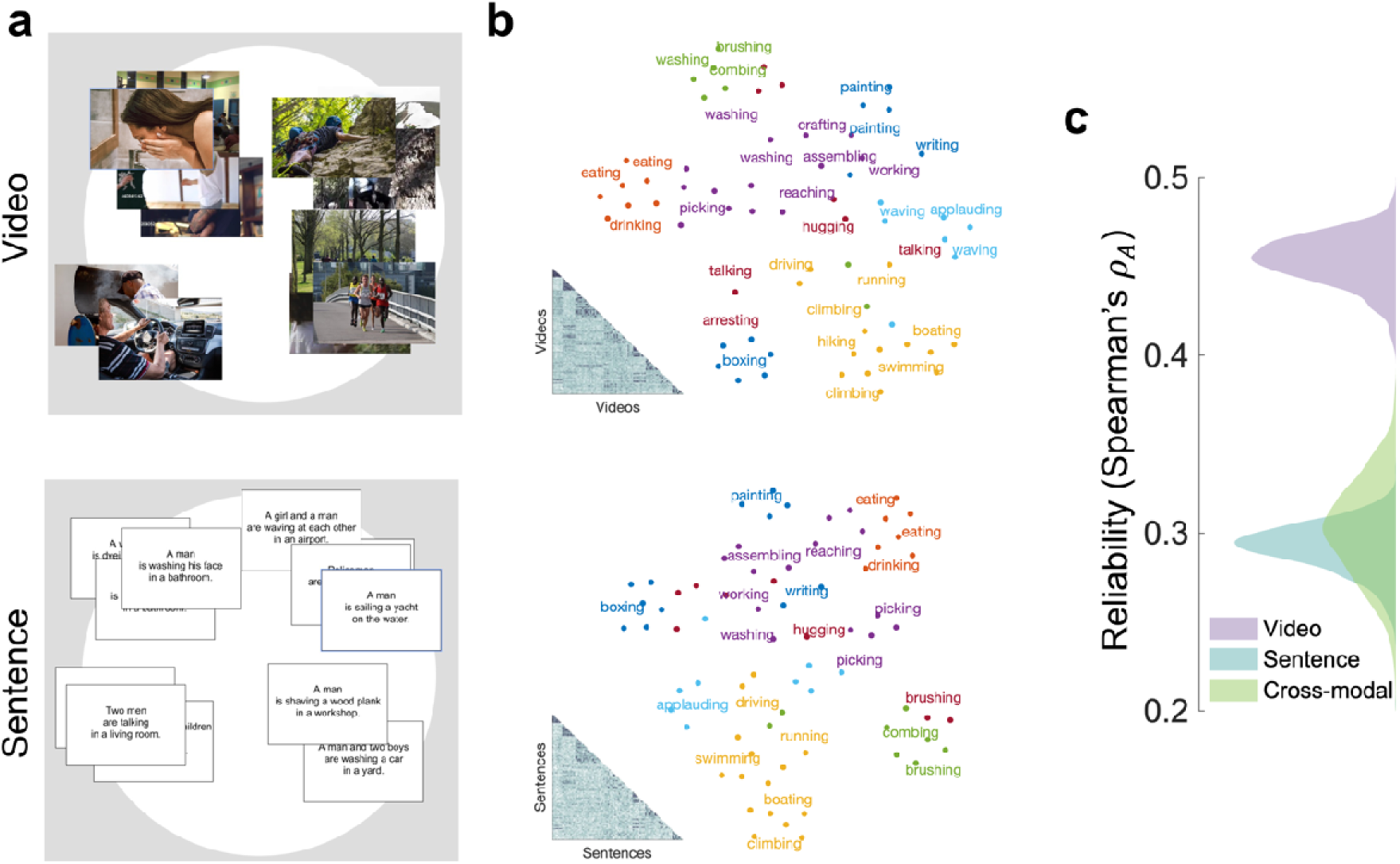
Multiple arrangement task and results. **a,** Examples of stimulus arrangements that could be mad by participants. The videos or sentences were presented around a circular arena. Participants arranged them inside the arena according to the actions’ semantic similarity. Images shown are public domain examples of the types of videos participants might see. **b,** Representational dissimilarity matrices (RDMs) were generated from eac participant’s arrangement. Here, the average RDM is shown in the lower-left corner, together with a t-distribute Stochastic Neighbor Embedding (tSNE) representation color-coded by action class and labeled with action verbs. **c,** Reliability of similarity judgments within and across modalities (correlation between subsets), pointing to a share structure across modalities. Distributions are based on 1000 data splits with N = 16 across all datasets.

The resulting dissimilarity matrices reflected a semantic organization when inspected using t-distributed Stochastic Neighbor Embedding (tSNE) plots (**Figure 2b**). The reliability of the data, quantified by subsampling two sets of 16 participants and correlating their averaged RDMs (1000 iterations), was higher in the video set (p_A_=0.45±0.01) than the sentence set (p_A_=0.29±0.01; Wilcoxon *z*=38.72, *p*<0.001). All reliabilities were significantly above chance (Wilcoxon *z*=27.39, *p*<0.001).

Across modalities, the average behavioral RDMs were substantially correlated ( =0.4). In a subsampling analysis drawing 16 participants from each modality 1000 times, the cross-modal correlation was =0.3±0.02, slightly higher than the internal reliability of the sentence dataset (Wilcoxon *z*=8.42, *p* <0.001; **Figure 2c**) and pointing at a strong cross-modal overlap.

### Features in action categorization

We first evaluated the contribution of each feature to behavior by correlating each feature RDM to each participant’s behavioral RDM (**Figure 3a**). Among the semantic features, the broadest (action target and class) correlated significantly with behavioral similarity across videos and sentences. In the video set, the everyday activity category also correlated with behavioral similarity, suggesting that this feature may be more salient in visual actions. None of the contextual and modality-specific features contributed to behavior, except for the visual effectors feature coding the body parts involved in each action.

**Figure 3.**
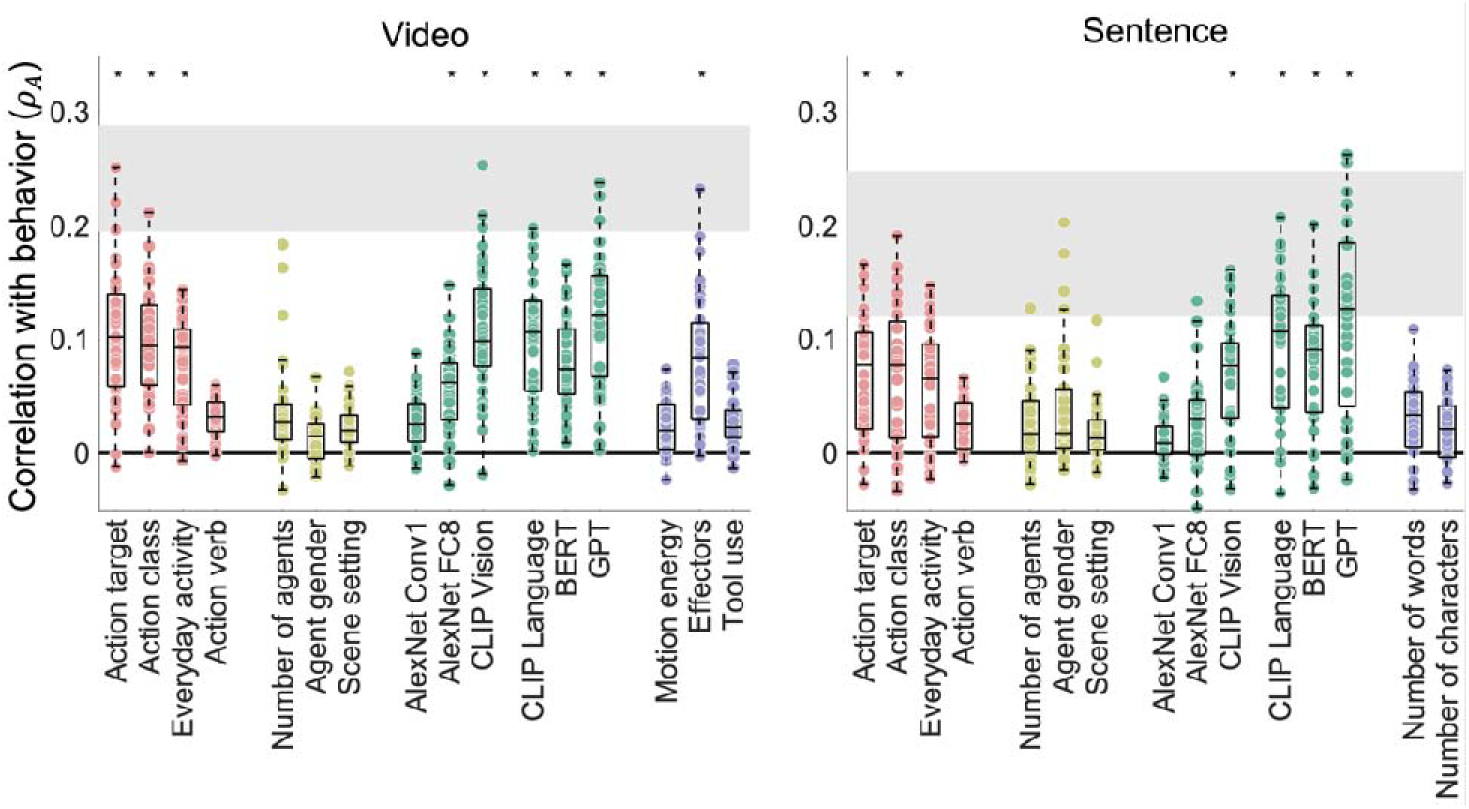
Broad semantic features and computational features best correlate with behavioral similarit within and across modalities. Individual feature-behavior correlations are plotted against the noise ceiling (in gray). Dots are individual subjects. Asterisks denote correlations significantly greater than zero (*p*<0.005).

Features extracted from computational models correlated with behavior acros modalities. In the video set, this included higher-level features from visual models (the last layer of AlexNet and CLIP’s vision module; see also **Supplementary Figure 1**), as well as all the language models (CLIP, BERT, and GPT). A model trained specifically for action recognition (ResNet-18 3D pretrained on Kinetics) did not correlate with behavior (**Supplementary Figure 1**).

In the sentence set, all LLM features, as well as visual features from CLIP, correlated with behavior, suggesting that these models represent conceptual information about actions. GPT model embeddings were the only feature to approach the noise ceiling in the sentence set, and also reached the highest correlation with behavior in the video set.

### Disentangling semantic features in action categorization

Even moderate correlations between related features (**Figure 1b**) could affect their contributions to the behavioral data, making their independent roles difficult to assess. We used cross-validated variance partitioning^12,55^ to evaluate the unique and shared variance in behavior predicted by different features. We trained and tested hierarchical regression models to predict similarity judgments within the video and sentence sets, as well as across modalities. Leave-one-out cross-validation was conducted across actions, with different participants’ data used for training and testing (see Methods: Variance partitioning).

First, to evaluate to what degree the behavioral datasets captured semantic information, we grouped our feature set into three categories: semantic (action target, class, activity, and verb); social (number of agents and agent gender); and perceptual (scene settings and selected modality-specific features; see Methods: Variance partitioning).

Together, the three groups of features approached or reached the noise ceiling (split-half data reliability) in predicting variance within and across modalities (**Figure 4a**). In all cases, semantic features explained more unique variance than the other groups (*p*<0.001, randomization testing). Social features explained a significant amount of unique variance in sentence similarity (*p*<0.001), but not video similarity (*p*=0.24) or across modalities (*p*=0.19). Perceptual features explained unique variance in video similarity (*p*<0.001), but not sentence similarity (*p*=0.15) or across modalities (*p*=0.31). This points to differences in the features aiding the semantic categorization of videos and sentences, and shows that only semantic features are shared across modalities.

**Figure 4.**
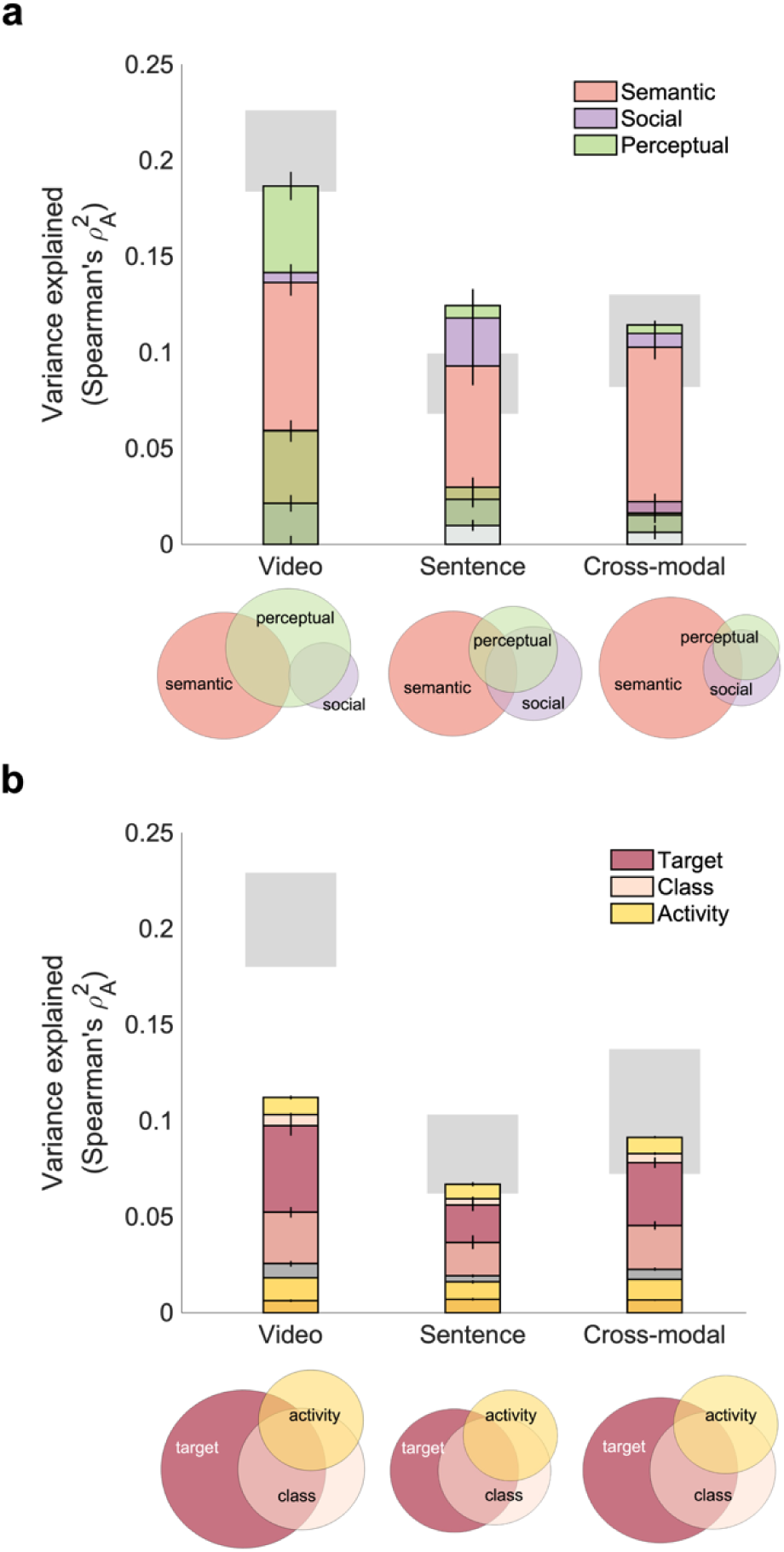
Variance partitioning: semantic features, particularly the action target and class, predict the most variance in behavior within and across modalities. **a,** Stacked plots and Venn diagrams showing the unique an shared contributions of semantic, social, and perceptual features to behavior (see Methods). Error bars are ±SD an the data reliability (min-max range, split-half _p_^2^) is shown in gray. As expected, semantic features explain the most variance. **b,** Unique and shared contributions of the three most predictive semantic features. The action target explains the most variance, including variance shared with the action class.

Next, we disentangled the broad semantic features (action target, action class, and everyday activity) that best correlated with behavior (**Figure 4b**). Both within and across modalities, the action target explained the most unique variance (*p*<0.001), as well as a large amount of variance shared with the action class (*p*<0.001). These results are in line with work suggesting that action goals are essential in action categorization^12,43^, and highlights the overlap between different action taxonomies.

### Large language model features predict human similarity judgments

Given how well GPT embeddings correlated with behavioral judgments in both modalities, we conducted an additional analysis to assess whether they shared information with the semantic features. To this end, we entered the GPT embedding RDM into a variance partitioning analysis together with the action target and class (**Figure 5a**).

**Figure 5.**
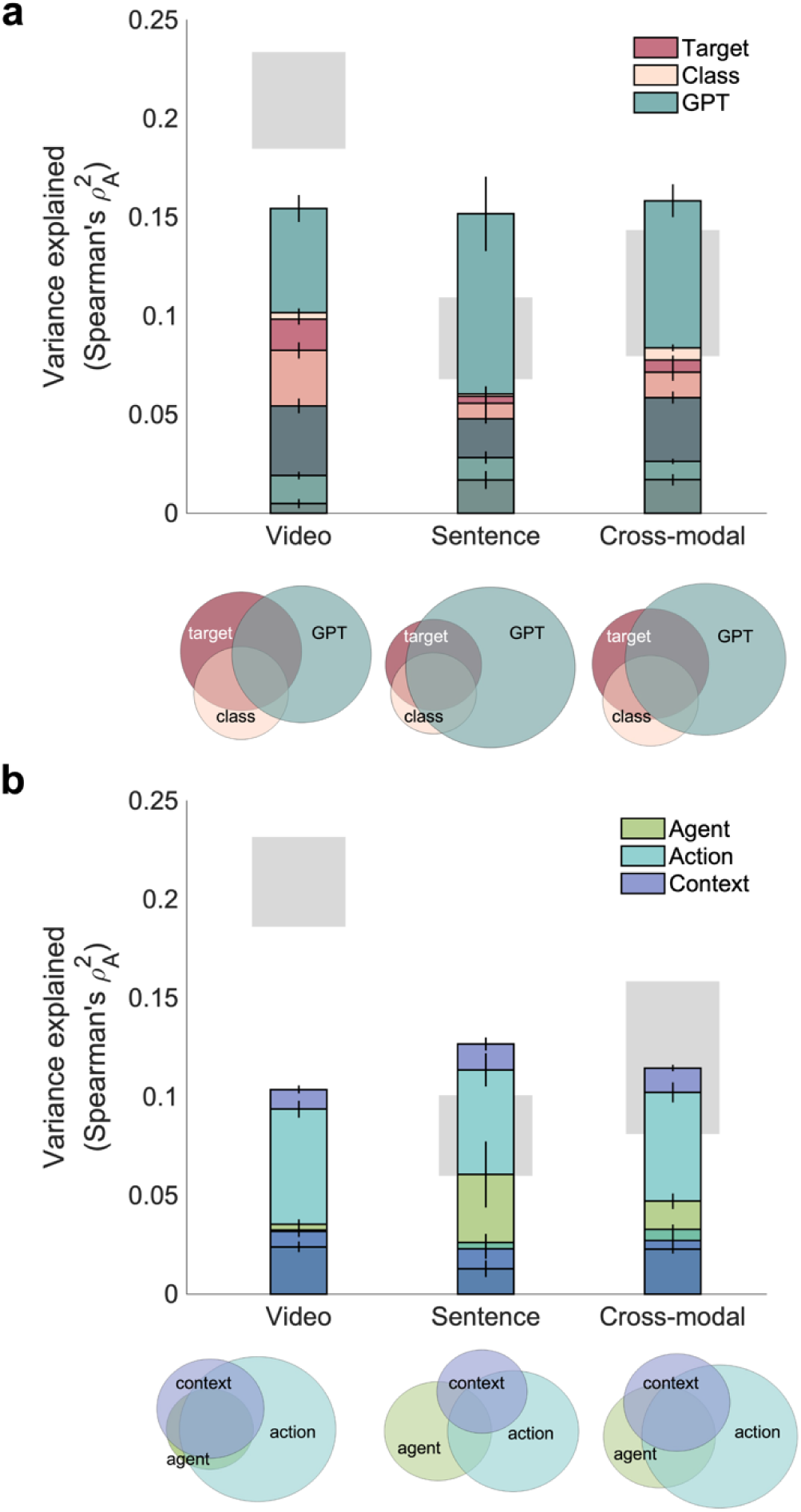
Variance partitioning: understanding GPT embeddings. **a,** Unique and shared variance i behavior explained by GPT embeddings, action target, and action class. GPT captures more, and different, variance in behavior than the two main semantic features, alongside shared variance with the action target and class. **b,** Unique and shared contributions of GPT embeddings based on the agent, action, and context parts of sentences. Most of the embeddings’ explanatory power reflects action information, as well as agent information when predicting sentence similarity.

Both within and across modalities, GPT uniquely explained a large portion of the variance in behavior (*p*<0.001), suggesting that GPT captures more information relevant to human categorization than individual semantic features. There was also significant shared variance between action target and class in the video dataset (*p*<0.001), and between action target and GPT embeddings in the video dataset and across modalities(*p*<0.001).

As naturalistic sentences contain rich information, it is difficult to know what features are captured by the sentence-level GPT embeddings. To assess this, we extracted separate GPT embeddings for the three sections of each sentence: the agent (e.g. *women*), the action (e.g. *are swimming*), and the context (e.g. *in a pool*). We used these embeddings to predict behavior in a variance partitioning analysis (**Figure 5b**), and found that action embeddings explained most of the unique variance both within and across modalities (*p*<0.001). However, in the sentence dataset, agent embeddings also contributed unique variance (*p*=0.001). Shared variance between all three features contributed significantly in the video dataset and across modalities (p<0.001) This suggests that that GPT embeddings capture both semantic and contextual information.

The hypothesis-driven semantic features were binary, allowing us to compare different levels of action categorization (see Methods: Feature definitions). However, this may have put them at a disadvantage when compared with the GPT embeddings, despite the use of a correlation metric that can handle differences in feature complexity. In a control analysis, we used FastText^78^, a pretrained word embedding, to generate semantic models based on text embeddings of the action target and class. This yielded similar results (**Supplementary Figure 2**), suggesting that the findings were not driven by differences in model complexity.

## Discussion

Using a naturalistic, multimodal dataset and semantic similarity judgments, we interrogated how action concepts are organized in the mind across vision and language. We reveal a shared organization along broad semantic axes reflecting the actions’ target and behaviorally relevant class. Furthermore, we show that language model embeddings accurately capture this shared semantic structure.

### A cross-modal organization of actions in the mind

Using a semantic similarity task and a naturalistic set of action videos and sentences, we found a shared semantic organization of actions in the mind across vision and language. This was reflected in the similar categorization of videos and sentences and the high reliability of similarity judgments across modalities (**Figure 1c**).

Previous behavioral studies have focused on either visual stimuli^11,14,15^ or text data^17,49^ to understand how actions are organized in the mind. However, neural investigations using both visual and text stimuli have found modality-invariant encoding of actions in the brain^25,26^. Our work links these findings to a shared representational space across vision and language.

Although semantic features explained most of the variance in behavior across modalities, the contribution of contextual features was different for videos and sentences (**Figure 4a**). In the video arrangement task, perceptual features (scene setting, motion energy, AlexNet final layer activations, tool and effector features) made a significant unique contribution, while social features (number and perceived gender of agents) explained little variance. Conversely, sentence similarity reflected social features, but not perceptual features, alongside semantic information. These differences open exciting avenues for identifying the building blocks of natural action categorization in vision and language.

Previous work has highlighted the important role of social features, particularly sociality (often defined as person-directedness), in intuitive visual action processing^8,9,43,56^. However, sociality can be difficult to disentangle from other semantic and perceptual features, such as action category and number of agents^8,11,57^. Here, the action target feature, which included aspects of both sociality (person-directedness) and transitivity (object-directedness), explained behavioral judgments best. In contrast, lower-level agent-related features did not capture much variance in the video behavioral data.

While efforts were made to curate a diverse action space representing many categories at different levels of abstraction, the choice of actions is likely to have a large impact on the behavioral similarity judgments. Furthermore, individual variability in arrangements placed limits on effect sizes and was reflected in the difference in reliability between visual and language stimuli. Here, we focused on determining shared, generalizable axes for action categorization in our dataset. Individual differences in categorization, and how these apply to different sets of actions, remain open questions for future research.

### Actions are categorized according to their target and class

Among semantic features, action target, action class, and activity category correlated best with the individual similarity judgments (**Figure 3a**). As these features were moderately correlated (particularly action target and class: p_A_ = 0.27), we performed a variance partitioning analysis to assess their unique and shared contributions (**Figure 4b**). Action target uniquely predicted behavioral judgments within and across modalities, in line with previous research highlighting the importance of action goals in organizing action representations in brain and behavior^12,32,58^. Action class, a feature inspired by a proposed human motor repertoire distinguishing between behaviorally relevant action categories^31,41,42^, shared variance with the action target. This likely reflects the fact that motor goals are generally shared by exemplars within each proposed action class^41^. It is interesting, however, that action class contributed little unique variance in behavior, suggesting that we should look to broader behaviorally relevant features as organizing axes in action understanding. Indeed, the primacy of goals can arguably allow for more flexible handling of the vast human action space.

Video similarity judgments also correlated with the effectors feature describing what body parts were involved in each action. This information, although highly diagnostic of action goals and types^9,33,41^, was not present in the sentence versions of the stimuli.

### Language models capture human action concepts

Recently, neural networks have made significant strides in predicting human behavioral judgments^59–62^ and neural responses to naturalistic stimuli across vision^63–68^ and language^69–72^. However, actions remain a difficult domain to tackle for even state-of-the-art algorithms. We assessed both computer vision models (AlexNet, CLIP, and others; see **Supplementary Figure 1**) and natural language processing models (CLIP, BERT, and GPT text embeddings) in their ability to capture the semantic organization of action similarity judgments. Features extracted from both types of models were used to explain behavioral data across modalities, to assess how well they capture the covariance between language and the visual world^73–76^.

Visual and textual features extracted from CLIP, a multimodal transformer trained on image and text pairs, captured behavioral representations well within and across modalities, but were surpassed by GPT embeddings (**Figures 3, 5**). GPT embeddings explained behavioral similarity better than all other language models, and better than individual semantic features. They captured video similarity better than vision models and uniquely explained a large portion of the variance both within and across modalities. They also shared significant variance with the action target, highlighting the importance of this feature in structuring action representations. While the combined semantic features (particularly action target and class) best modeled the modality-invariant component in the behavioral data, the ability of GPT embeddings to explain variance both within and across modalities suggests an advantage related to their multidimensionality. Furthermore, the ability of LLM features to capture semantic representations shared across vision and language points to the extraction of semantic information as an important goal in human vision^77^.

Furthermore, an analysis comparing GPT embeddings extracted from the agent, action, and context parts of each sentence shows that most of their explanatory power originated from the action itself (**Figure 5b**). While a binary verb model did not strongly correlate with behavior, GPT embeddings based on verbs capture complex relationships that also reflect higher-level features. In the video dataset, context-related GPT embeddings did not capture perceptual variance as successfully as image features (**Figure 4a**), and thus did not explain all the variance in similarity. However, in the sentence dataset, both action and agent embeddings explained significant variance. Together, the GPT embeddings reached the noise ceiling in both sentence and cross-modal prediction, pointing at the strength of this model in capturing the conceptual organization of actions. Collectively, these results demonstrate the generalizability of text representations used by LLMs across tasks and modalities, as well as the potential for using human behavioral data and insights to better understand these models.

### Features and categories in minds and machines

Since the discovery of functional selectivity for object categories in the human brain^79–81^, a growing body of work has investigated the role of visual features in explaining or supporting categorization^61,82–84^. In action perception, evidence for categorical, invariant representations^1,6,11,28,85,86^ has grown alongside evidence for featural representations along the ventral and dorsal streams: from bodies and their motion^81,87–91^ to information about tools and effectors^2,27,33,43^ and complex social information^8,9,41,92–94^. Importantly, the use of naturalistic stimuli has been essential in uncovering these rich representations, adding to growing evidence of the impact of context in visual processing^37,95–98^, and pointing to the importance of actions as a step towards natural event understanding.

Here, we focused on categorical models of semantic organization, grouping actions into classes at different levels of abstraction or according to their targets, effectors, or the involvement of tools. Our results show that a broad categorical model (i.e., whether an action is directed towards a person, an object, or the self) determines much of the semantic organization of actions in the mind, which is consistent across vision and language. However, it remains to be seen whether this organizing feature should be thought of as truly categorical or continuous, mapping actions along a spectrum according to whether they aim to instill change in the self, the environment, or other people.

Although the semantic similarity task ensured a focus on actions, our stimuli were naturalistic by design, involving interactions between agents and objects in a variety of contexts. Thus, perceptual and social features contributed to the similarity judgments, highlighting the role of actions as a bridge between visual domains and an essential component in event understanding. Action goals in particular are thought to play an important part in the perception and segmentation of events^99^. These results thus open opening exciting avenues towards understanding how natural events are processed and organized in the mind.

As artificial intelligence models make strides towards human-like performance across tasks and domains, new questions arise about their internal representations and their alignment with the human mind. Our results highlight the ability of language model embeddings to capture human-like action representations, adding to recent work that suggests a key role for cognitive science in measuring and narrowing the gap between model and human representations^62,100–102^.

Together, our results characterize the semantic organization of naturalistic actions across vision and language, and open exciting avenues for investigating conceptual representations and their building blocks in minds and machines.

## Methods

### Stimuli

#### Videos

The stimulus set consisted of 95 short videos and sentences depicting everyday actions. Actions were defined at four levels of abstraction from specific (action verb) to broad (action target; **Figure 1a**; see Methods: Feature definitions).

Videos were selected from the Moments in Time dataset^44^. First, we inspected the Moments in Time verb labels to identify actions belonging to eight main classes (defense, ingestion, locomotion, manipulation, self-directed actions, gestures, social interactions, and symbolic actions)^41,42^. We then curated an initial set of 249 videos with the aim of representing each class through a variety of verbs and everyday activities (see Methods: Feature definitions). Videos were chosen that were horizontally oriented (landscape), shot in color, and of reasonable image quality. Each video depicted a single, continuous human action. We excluded videos where agents interacted with the camera. The selected videos varied along multiple axes related to the action (action target, tool use), agents (number, gender), and scene setting, which were manually labeled by the experimenters across the video set.

Since features can covary in naturalistic stimulus sets, we performed further stimulus selection to reduce feature correlations. To this end, we randomly drew 10,000 samples of 100 videos from the initial stimulus set. Each sample included 10 videos per action class, except for locomotion and manipulation, which were represented by 20 videos each to reflect their action diversity.

Out of all samples, we selected the set of stimuli with the lowest overall correlation across a range of visual, semantic, agent and action features (mean Spearman’s p=0.07, SD=0.13). We manually adjusted this set to remove videos that were too visually similar to others in the same category. The final set consisted of 95 videos, with 7-20 videos per action class.

All videos were resized to 600 x 400 pixels. To achieve this, we first applied the *imresize* function in MATLAB R2021b using bicubic interpolation. Next, the videos were cropped to the correct aspect ratio either centrally or using manually determined coordinates to ensure the action remained clearly visible. Using the *videoWriter* object in MATLAB R2021b, the videos were trimmed to 2 s centered around the action and resampled to 20 fps.

#### Sentences

Sentences were generated to succintly describe the action in each video, using a consistent format: Agent + Action [+ Object, if present] + Context; for example: *A girl and a boy are eating burgers in a kitchen. Two men are talking in a living room.* (see **Appendix 1** for the full list of sentences).

The number and perceived gender of agents were specified where clearly visible. Actions were accompanied by their objects where applicable. There was no difference in overall length between sentences that included an object and those that did not (*t*(82.7)=1.22, *p*=0.22). The context was described using a single word or phrase. The most specific wording possible was used to reflect the level of detail present in the video. Although detail was necessarily lost in the transition between modalities, we emphasized natural phrasing and included information about agents, actions, objects, and contexts to ensure a clear one-to-one correspondence between actions and videos.

Sentence stimuli were 400×600 pixel images with gray borders, ensuring visual consistency with the video stimuli (**Figure 2a**). Each image displayed a sentence in black Arial font on a white background. For readability, each sentence was split into three lines corresponding to the three sentence components (agent, action, and context).

### Participants

A sample of 41 participants completed the video arrangement experiment (39 after exclusions, mean age 18.08, SD 0.84, 22 female, 16 male, 1 non-binary). A different cohort of 37 participants completed the sentence arrangement experiment (32 after exclusions, mean age 18.16, SD 0.45, 19 female, 13 male). Participants were recruited through the Department of Psychology Research Participation Pool at Western University. All procedures for online data collection were approved by the Western University Research Ethics Board, and informed consent was obtained from all participants.

### Multiple arrangement

We conducted two multiple arrangement experiments to understand semantic action representations across vision (Experiment 1, using videos) and language (Experiment 2, using sentences).

The experiments were conducted on the Meadows platform (https://meadows-research.com/). Participants were instructed to arrange the videos or sentences according to the actions’ semantic similarity, placing similar actions closer together and different actions further apart.

The task took 90 minutes to complete and an average of 186 (videos) or 215 (sentences) trials. The difference between the numbers of trials in the two tasks was not significant (*t*(60)=0.97, *p*=0.34). Each trial started with a maximum of 12 videos or sentences arranged around a circular arena. The videos played automatically on hover. All stimuli had to be dragged-and-dropped inside the arena before the trial ended. Participants viewed different subsets of stimuli during each trial. Stimulus selection was guided by a ‘lift-the-weakest’ algorithm so as to optimize the signal-to-noise ratio of the distance estimates^54^.

Before the main task, participants completed a training trial in which they arranged the same three videos (in Experiment 1) or sentences (in Experiment 2). Participants were excluded if their training data correlated poorly to the average of all other participants’ data (over 2 SDs below the mean). Participants were also excluded if they responded incorrectly to a catch trial requiring them to label the action in previously seen stimuli.

Using inverse MDS, participants’ arrangements were used to generate behavioral dissimilarity matrices quantifying the Euclidean distances between all pairs of videos. We visualized the clustering of action categories using t-SNE representations of the average RDMs, which display the stimuli as points in a 2D space.

### Feature definitions

#### Semantic features

Actions can be defined at several levels of abstraction, from specific verbs to broad classes. Which of these guides human judgments about the meaning of actions? We tested several features to understand the structure of semantic action representations in the mind. Each feature was converted into a representational dissimilarity matrix (RDM) quantifying the dissimilarity between stimuli along that axis.

Our four main hypothesis-driven semantic features were: the action target (whether an action was directed towards an object, a person, or the self); action class (based on a putative human motor repertoire^41^); everyday activity (based on activity categories from the American Time Use Survey^40^); and action verb (based on the Moments in Time verb labels; see **Figure 1a** for a complete list of semantic categories). These were binary RDMs, in which distances between stimuli belonging to the same category were set to 0, and distances between stimuli belonging to different categories were set to 1.

#### Contextual features

To capture information about the agents (social features), we included the number of agents (labeled on a three-point scale from 1, one agent present, to 3, three or more agents present) and their perceived genders. To capture action context, we included scene setting as a perceptual feature (indoors/outdoors). Contextual features were labeled by experimenters and converted to binary or Euclidean distance RDMs.

#### Computational features

To assess whether computational models can capture human semantic judgments, we extracted activations from computational models trained on visual and text stimuli.

As the videos were short and did not contain major visual changes across frames, we extracted most activations using the first frame of each video. We used the AlexNet feedforward neural network pretrained on ImageNet, and extracted features from its first convolutional layer (Conv1, to capture low-level visual properties), and its last fully connected layer (FC8, to capture higher-level features). We also extracted features from CLIP, a multimodal transformer trained on publicly available image-text pairs^51^. As CLIP contains both image and text encoders, we used it to extract features from both videos and sentences. AlexNet features were extracted using the Deep Learning toolbox in MATLAB R2022a, while CLIP features were extracted using the THINGSvision^103^ toolbox in Python 3.10.9. Additional model features, including ResNet-50, VGG-16, CORnet-S, CORnet-RT (all pretrained on ImageNet), and ResNet-18 3D (pretrained on Kinetics) were extracted using the THINGSvision toolbox or PyTorch and the *torchextractor* package in Python 3.10.9 (Supplementary Figure 1).

We also extracted embeddings (features measuring the relatedness of text strings) from Google’s BERT^52^ transformer and OpenAI’s embedding model associated with its GPT^51,53^ transformers. We used the latest version of GPT embeddings at the time of writing, *text-embedding-ada-002*, which outperform previous OpenAI embedding models despite being lower-dimensional. To assess the contributions of different sentence components, we also extracted separate embeddings for the agent, action, and context portion of each sentence. All text features were extracted from the Hugging Face transformers library^104^ using PyTorch in Python 3.10.9. RDMs were created using pairwise Euclidean distances between feature vectors.

#### Modality-specific features

Finally, we quantified modality-specific features to capture additional sources of variance. For videos, these included motion energy computed using a pyramid of spatio-temporal Gabor filters as implemented in the *pymoten* package^105^. We also included two video-specific action features which were not present or discernible in the sentences: tool use (binary feature: whether the action was tool-mediated), and effectors (binary vectors indicating the body parts involved in each action: face/head, hands, arms, legs, and torso).

For the sentence set, we created two low-level features reflecting the number of words and characters in each sentence.

### Representational Similarity Analysis (RSA)

To assess each feature’s contribution to the human similarity judgments, we calculated Spearman’s p_A_ correlations between each feature RDM and each participant’s behavioral RDM. While Spearman’s p can favour categorical models with many tied ranks, Spearman’s p_A_ solves this issue using random tiebreaking to compute the expected correlation^106^. All RDMs were normalized using min-max normalization.

We assessed the significance of the correlations using sign permutation testing (5000 iterations, one-tailed). P-values were omnibus-corrected for multiple comparisons using a maximum correlation threshold across models^107,108^.

A noise ceiling was calculated by correlating each participant’s RDM to the average RDM across participants (upper bound), as well as to the average RDM excluding the left-out subject (lower bound)^109^.

### Variance partitioning

To disentangle the contributions of specific sets of features to the behavioral similarity judgments, including their unique and shared contributions, we performed cross-validated variance partitioning within and across modalities.

We first evaluated whether the behavioral data reflected semantic information more than social and perceptual information. To this end, we selected three groups of features: semantic (action target, action class, everyday activity, and action verb); social (number of agents and agent gender); and perceptual (for videos: scene setting, tool use, effectors, motion energy, and features from the FC8 layer of AlexNet; for sentences: scene setting and number of words; cross-modal: scene setting). Each group of features was considered a separate predictor in this analysis.

We repeated this analysis with different sets of predictors to evaluate: (1) the variance explained by each semantic feature that significantly correlated with the behavioral similarity data (action target, action class, and everyday activity); (2) the variance explained by GPT embeddings compared to the most predictive semantic features (action target and action class); (3) the variance explained by GPT embeddings separately extracted from the agent, action, and context portion of each sentence.

At each iteration, we fit seven regression models using every possible subset of the three predictors and evaluated them using leave-one-out cross-validation. Across 95 iterations, the models were fit using the data from 94 out of the 95 actions (4371 pairs). We then predicted the similarity judgments for the held-out data (i.e., the dissimilarities between the held-out action and all others, amounting to 94 pairs). To assess how well the models predicted the held-out data, we calculated the squared Spearman’s p_A_ correlation between the predicted and actual similarity judgments, as an approximation of the amount of variance explained. We then used this to calculate the unique and shared variance contributed by each group of features^8,55,110^.

Importantly, the training and test sets were drawn from different participants. In the within-modality analyses, we performed 100 random splits of each behavioral dataset into training and test sets of approximately equal sizes, which were averaged before performing leave-one-action-out cross-validation as described above. Results were averaged across 95 iterations of leave-one-out cross-validation and 100 dataset splits.

In the cross-modal analysis, we drew 100 random subsets of 20 participants from each behavioral dataset. At each cross-validation iteration, we used video data for training and correlated the predicted responses with the sentence data (and vice versa). We averaged results across both directions of training and testing.

All results were tested against chance using one-tailed sign permutation testing (5000 iterations, omnibus-corrected for multiple comparisons).

## Acknowledgements

The authors would like to thank Leyla Isik for helpful comments on the manuscript and Martin Hebart for code calculating the Spearman’s p_A_. This material is based upon work supported by BrainsCAN at Western University through the Canada First Research Excellence Fund (CFREF) to Y.M., a Western Interdisciplinary Development Initiatives Grant to Y.M. and J.C.C., a Vector Institute for Artificial Intelligence Research Grant to Y.M. and D.C.D., and a Western Postdoctoral Fellowship to D.C.D.

## Data availability

The data and results have been archived as an Open Science Framework repository (https://osf.io/c5bdv/).

## Code availability

Analysis code is available on GitHub (https://github.com/dianadima/multi-action).

## Notes

### Competing Interest Statement

The authors have declared no competing interest.

### Summary of Updates

Updated methodology, edited for clarity, and updated Supplementary Information

