## Supplementary for "Shared representations of human actions across vision and language"

### Supplementary Information

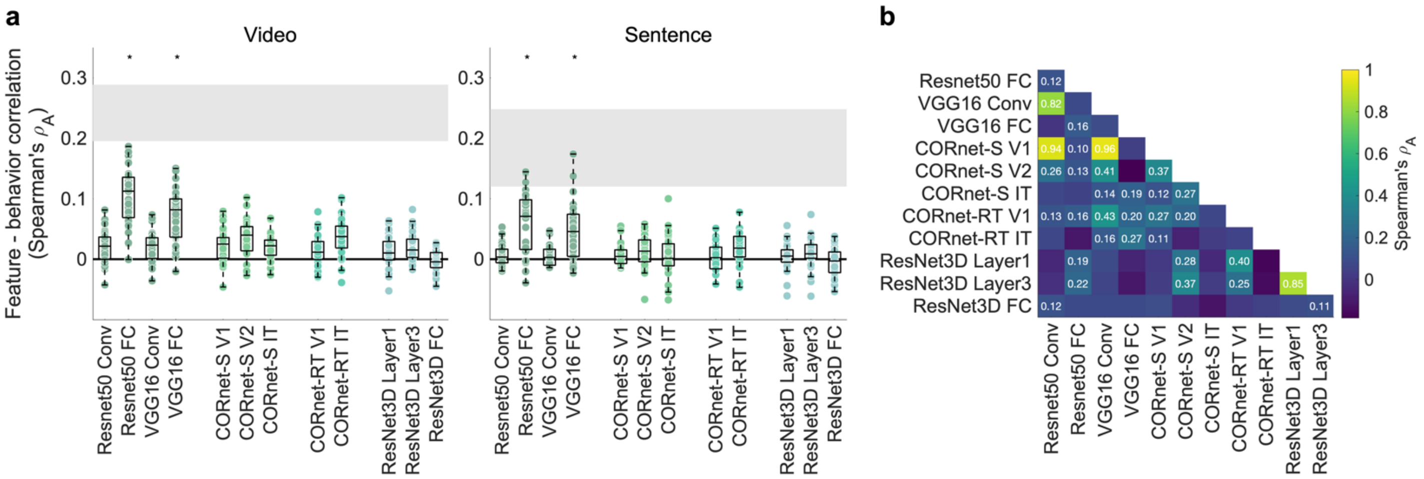


**Supplementary Figure 1**. Correlations between the behavioral similarity judgments and video features extracted from ResNet-50, VGG-16, CORnet-S, the recurrent CORnet-RT (all pretrained on ImageNet) and ResNet-18 3D (pretrained on Kinetics). **a,** Only the fully connected layers of ResNet-50 and VGG-16 significantly correlate with behavior ($\alpha$=0.005). Although ResNet-18 3D was trained for action recognition, it does not reflect human action categorization behavior. **b,** The correlations between computational model RDMs show significant overlap in the features extracted by different models, with the notable exception of the Kinetics-trained ResNet-18 3D.

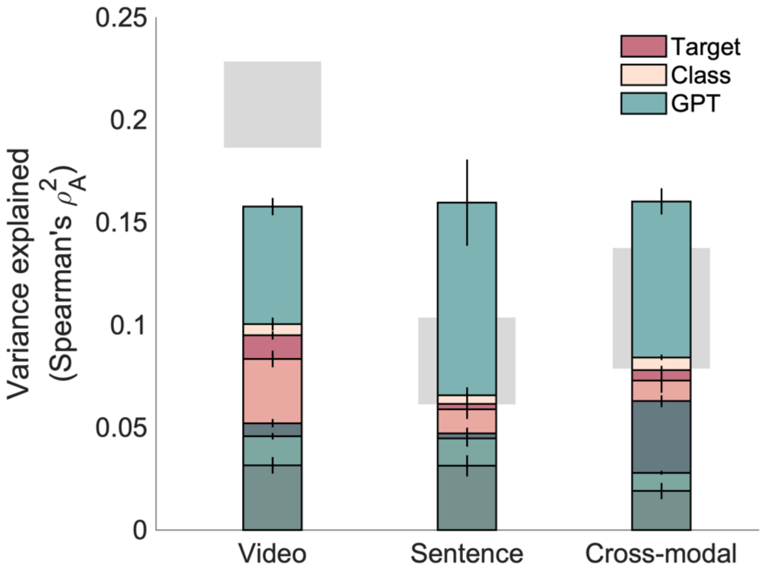


**Supplementary Figure 2**. GPT sentence embeddings explain more unique variance than the semantic features, even when using text embeddings to generate semantic feature RDMs. FastText word embeddings were generated for each everyday activity, action class, and action target, and semantic RDMs were obtained by calculating the pairwise distances between the 300-dimensional vectors.

### Appendix 1: List of sentences

Two women are boxing in a gym.

A woman and a man are practising defense in an attic.

Two men are fighting in the street.

Two men are fighting in a ring.

Two athletes are fencing in a gym.

Two men are boxing outdoors.

Two men are dueling in a park.

A girl is eating cereal at a table.

A man is drinking beer in a bar.

A man is drinking coffee on a balcony.

A man and a woman are eating breakfast on a balcony.

A girl and a boy are eating burgers in a kitchen.

A man and a woman are drinking coffee on a porch.

A man is eating a sandwich in a park.

A woman is sipping a cocktail by the sea.

A boy and a girl are eating ice cream in a park.

A group is eating watermelon on a lake shore.

A person is swimming in a river.

A boy is climbing a tree outdoors.

A man is rope climbing a cliff in the mountains.

A person is rappelling down a cliff.

A man and a woman are travelling in a car.

A woman is climbing an indoor climbing wall.

A person is biking over river rocks.

Two people are sailing a boat on water.

Two women are hiking a trail.

Children are running down a corridor.

A man is sailing a yacht on the water.

People are walking down a hallway.

People are hiking in the mountains.

A man is driving a car down a street.

A group is cycling on a street.

Women are swimming in a pool.

A woman is canoeing on a lake.

A man is hiking through snow.

A woman is climbing up the stairs in a lobby.

A woman is driving a car down a street.

Children are making a toy robot in a yard.

A baby is reaching for a phone in his crib.

A woman and a girl are rolling dough on a table.

A woman is picking up a seashell on a beach.

A man is shaving a wood plank in a workshop.

A worker is welding in a factory.

A girl is reaching for a plate in a kitchen.

A man is picking meat in a supermarket.

Two women are picking earrings in a shop.

A man is picking bananas in a supermarket.

A man and two boys are washing a car in a yard.

A man is mopping the floor in a shop.

A man and a boy are assembling a table in a room.

A woman and a man are picking apples from a tree.

A woman and a man are grilling on a terrace.

Men are barbecuing at a street stall.

Men are moving parcels in a post office.

A woman is drying dishes in a kitchen.

A girl is picking blackberries outdoors.

A man is watering plants in a garden.

A man is stretching on a bridge.

A woman is dyeing her hair in a bathroom.

A woman is brushing her teeth in a bathroom.

A man is brushing his hair in a bathroom.

A woman is combing her hair in a bathroom.

A man is drying his face in a bathroom.

A woman and two children are stretching in a dance hall.

A man is washing his face in a bathroom.

A woman is washing her face in river water.

A boy is brushing his teeth in a bathroom.

A woman is applauding in a park.

A woman is waving on a beach.

A woman is beckoning another in an office.

People are applauding in a restaurant.

People are waving at an outdoors concert.

A girl and a man are waving at each other in an airport.

Two people are waving on a ski piste.

An audience is applauding in a concert hall.

An audience is applauding in an auditorium.

Two girls are applauding at a party.

A groom and a bride are kissing in a park.

Two men are talking in a living room.

A woman is brushing a girl's hair on a balcony.

A man is lecturing students in a classroom.

A man and a woman are hugging in a coffeeshop.

A man is brushing another's hair in a barbershop.

A man is holding a baby in a living room.

Policemen are arresting a man on a street.

A woman is blow-drying a girl's hair in a bathroom.

A man and a woman are arguing in a room.

A woman is painting in a studio.

Three children are writing at a table.

A woman and two children are painting in a living room.

A man is writing in an office.

A man is spray painting a wall outdoors.

A woman is sculpting a statue in a studio.

A woman is painting a wall outdoors.

A woman is writing at a desk.
